# Food or physics: plankton communities structured across Gulf of Alaska eddies

**DOI:** 10.1101/2021.10.15.464534

**Authors:** Caitlin Kroeger, Chelle Gentemann, Marisol García-Reyes, Sonia Batten, William Sydeman

## Abstract

Oceanic features, such as mesoscale eddies that entrap and transport water masses, create heterogeneous seascapes to which biological communities may respond. To date, however, our understanding of how internal eddy dynamics influence plankton community structuring is limited by sparse sampling of eddies and their associated biotic communities. In this paper, we used 10 years of archived Continuous Plankton Recorder (CPR) data (2002-2013) associated with 9 mesoscale eddies in the Northeast Pacific/Gulf of Alaska to test the hypothesis that eddy origin and rotational direction determines the structure and dynamics of entrained plankton communities. Using generalized additive models and accounting for confounding factors (e.g., timing of sampling), we found peak diatom abundance within both cyclonic and anticyclonic eddies near the eddy edge. Zooplankton abundances, however, varied with distance to the eddy center/edge by rotational type and eddy life stage, and differed by taxonomic group. For example, the greatest abundance of small copepods was found near the center of anticyclonic eddies during eddy maturation and decay, but near the edge of cyclonic eddies during eddy formation and intensification. Distributions of copepod abundances across eddy surfaces were not mediated by phytoplankton distribution. Our results therefore suggest that physical mechanisms such as internal eddy dynamics exert a direct impact on the structure of zooplankton communities rather than indirect mechanisms involving potential food resources.

## Introduction

Oceanographic features such as flowing currents and rotating mesoscale eddies (approx. 100– 200 km wide) can horizontally transport water masses across ocean basins, creating heterogeneous seascapes that biological communities thrive on. Eddies, in particular, are spawned from instabilities in water density gradients or spun off from currents interacting with topography [1]. Due to entrapment of water at formation, eddies may contain water masses that are different from their surroundings and highly variable in composition (i.e., nutrients, salinity, temperature) [2,3]. For example, anticyclonic mesoscale eddies originating from the Alaska Current are known to transport warm, low salinity, nutrient-rich water containing coastal plankton offshore into the Gulf of Alaska [4,5]. In addition to horizontally transporting captured water from their origins, eddies vertically transport water throughout their lifetime (days to years). Generally, anticyclonic eddies downwell water through their core, whereas cyclonic eddies upwell deeper, colder, nutrient-rich water layers through their core to the sea surface [6–9]. This “eddy pumping” of nutrients primarily occurs during formation but the dynamics can become complicated by other processes such as perturbations to circulation or eddy-induced Ekman upwelling from wind stress that generally causes surface upwelling (downwelling) in anticyclonic (cyclonic) eddy centers [6,10], particularly in strong wind regions and seasons where surface currents are most affected.

Eddy-associated surface chlorophyll concentrations and spatial distributions are differently influenced over time by pumping mechanisms as eddies evolve from formation to maturation to decay [11]. Moreover, relaxation during decay leads to opposing vertical pumping mechanics from that of formation, with downwelling (upwelling) predicted at cyclonic (anticyclonic) eddy centers [12]. These processes, in combination with mechanisms such as eddy-induced shifts in the mixed layer depth and peripheral stirring with surrounding waters as the eddy travels [11,13], lead to complex combinations of eddy dynamics throughout the eddy lifespan and ultimately contribute to diverse biological communities in an otherwise oligotrophic basin.

Biological communities depend on eddy-driven transport, upwelling, and mixing of nutrient-rich water [14]. In the Gulf of Alaska, the growth of primary producers such as phytoplankton is often limited by minerals (e.g., iron [15]) that are more abundant in coastal waters or subsurface water layers, and these minerals can be reintroduced to depleted regions by the water transport and/or mixing mechanisms of eddies [5,16,17]. Eddy-driven vertical and horizontal advection of nutrients has bottom-up effects on biological communities: organisms from plankton to fish to top predators (i.e., seabirds and marine mammals) depend on such processes, and are often found in close association with eddies [4,18,19]. However, our understanding of biotic community structuring within eddies is limited, as it comes from research where only one to a few eddies were sampled *in situ* (e.g., [20–23]). Plankton sampling typically occurs by net tows at point locations that are biased toward hard-carapace organisms; by continuous capture via gauze mesh (i.e., Continuous Plankton Recorder or CPR) that captures soft bodied organisms, but is analyzed at large internals (18 km segments every 74 km); or by continuous optical imaging of small water volumes in single eddies [22,24]. Sampling few eddies or at large intervals is problematic as there is considerable variability in plankton communities within and across mesoscale eddies, which can be missed with coarse or sparse sampling. Increased sample sizes of eddies and plankton abundances within eddies enhances our ability to distinguish the physical mechanisms that are important in shaping biological communities across regions and eddy life stages.

Using 10 years of archived CPR data paired with satellite measures of various long-lived mesoscale eddies, we increased the temporal and spatial resolution of plankton sampled across eddies to understand the importance of differing eddy physical dynamics in driving the structure of biological communities (phytoplankton and zooplankton) in the Gulf of Alaska. With these data, we tested the following hypotheses: (1) the composition and organization of plankton communities across marine mesoscale eddies depends on rotational type and eddy origin; (2) the influence of physical eddy dynamics (e.g., eddy pumping and advection) on plankton community organization is greater in the early life stage; and (3) eddy dynamics influence zooplankton communities indirectly through their effects on phytoplankton. We used generalized additive models (GAMs) to explore the effect of sampling location relative to the eddy edge on plankton species abundances, while taking into account the effects of year, season, and time of day. We then implemented nonlinear path analysis to explore the direct and indirect biophysical relationships between eddy characteristics and plankton densities. These biophysical connections will provide insight into similar regions that may be lacking in *in situ* plankton sampling, but have satellite coverage. Moreover, a more nuanced understanding of biophysical relationships can lead to improved ecological forecasting of fish (e.g., salmonids) and other wildlife (e.g., whales and seabirds) of societal value that rely on this region of the North Pacific as feeding grounds.

## Methods

### CPR data

Archived CPR samples from transects across the Gulf of Alaska were analyzed according to standard CPR protocols [25,26], from the Strait of Juan de Fuca to Cook Inlet or the Aleutian Islands (Fig 1 – map of CPR sample locations/study area), spanning the months of March – October and December from 2002 to 2013 (Table 1). The CPR is towed behind the ship at a depth of 7 m, where water flows through the CPR entrance (1.27 cm^2^) into a continuous 270-μm silk mesh net that filters plankton. Phytoplankton and zooplankton captured in the CPR mesh were cut in 18.5-km segments and examined with microscopy to identify and count taxonomic groups. The date, time, and coordinates (degrees latitude and longitude) were associated with the midpoint of each segment. The phytoplankton taxonomic groups in this study included diatoms and dinoflagellates. The zooplankton taxonomic groups in this study included euphausiids, large copepods, small copepods, pteropods, hyperiids, chaetognaths, larvaceans, and microzooplankton. See [25] for further details on CPR methodology.

**Table 1.**
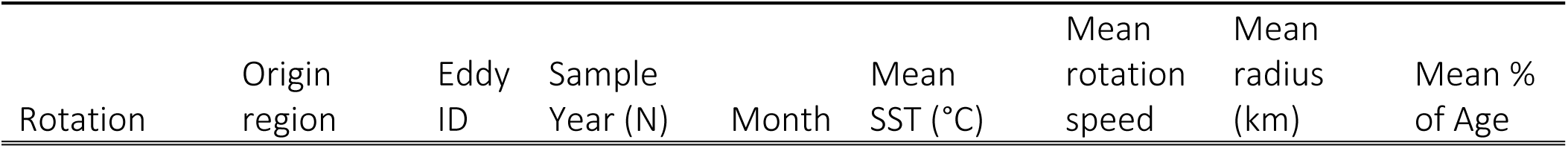
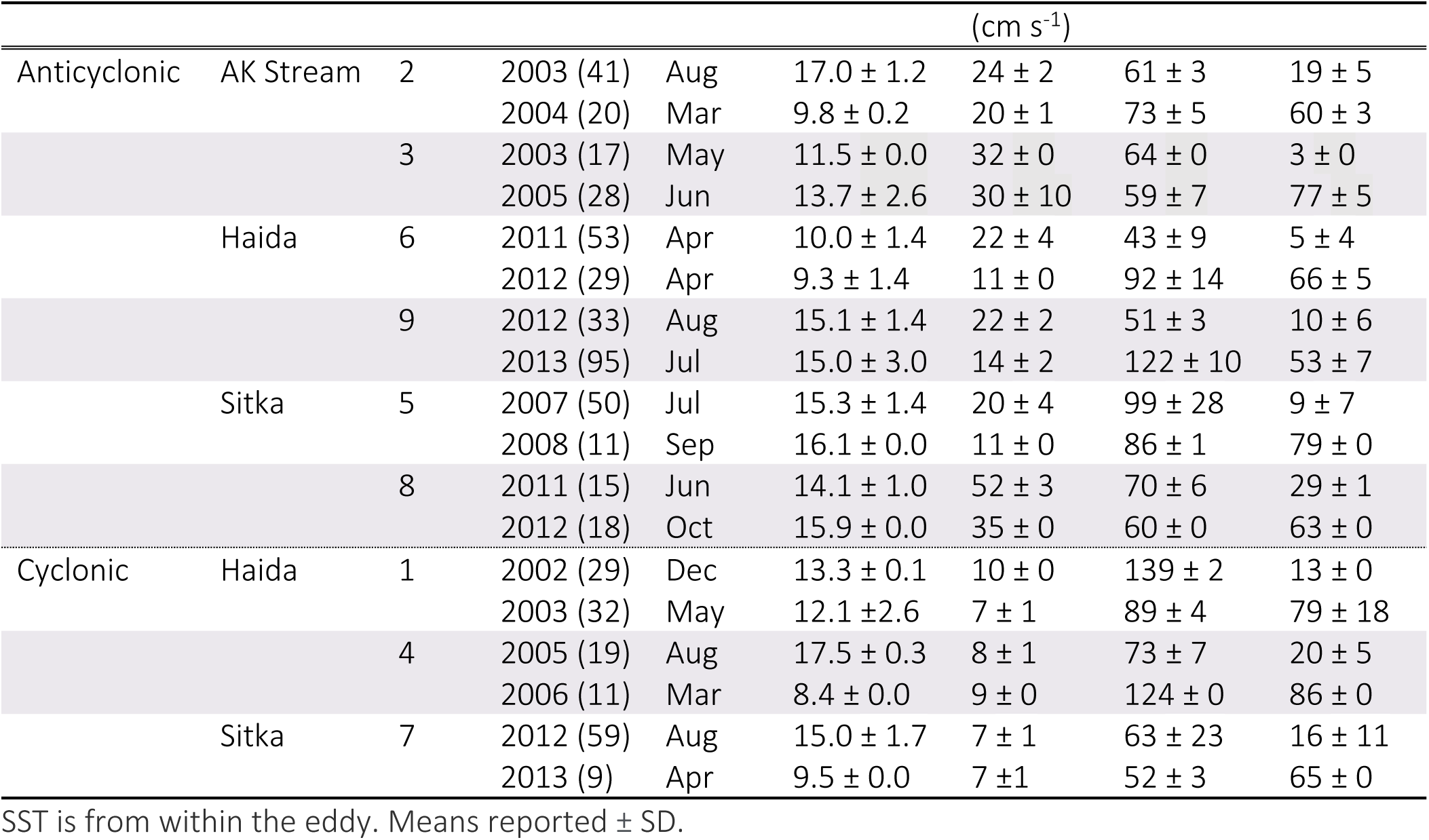
Summarized eddy characteristics during sampling periods.

**Fig 1.**
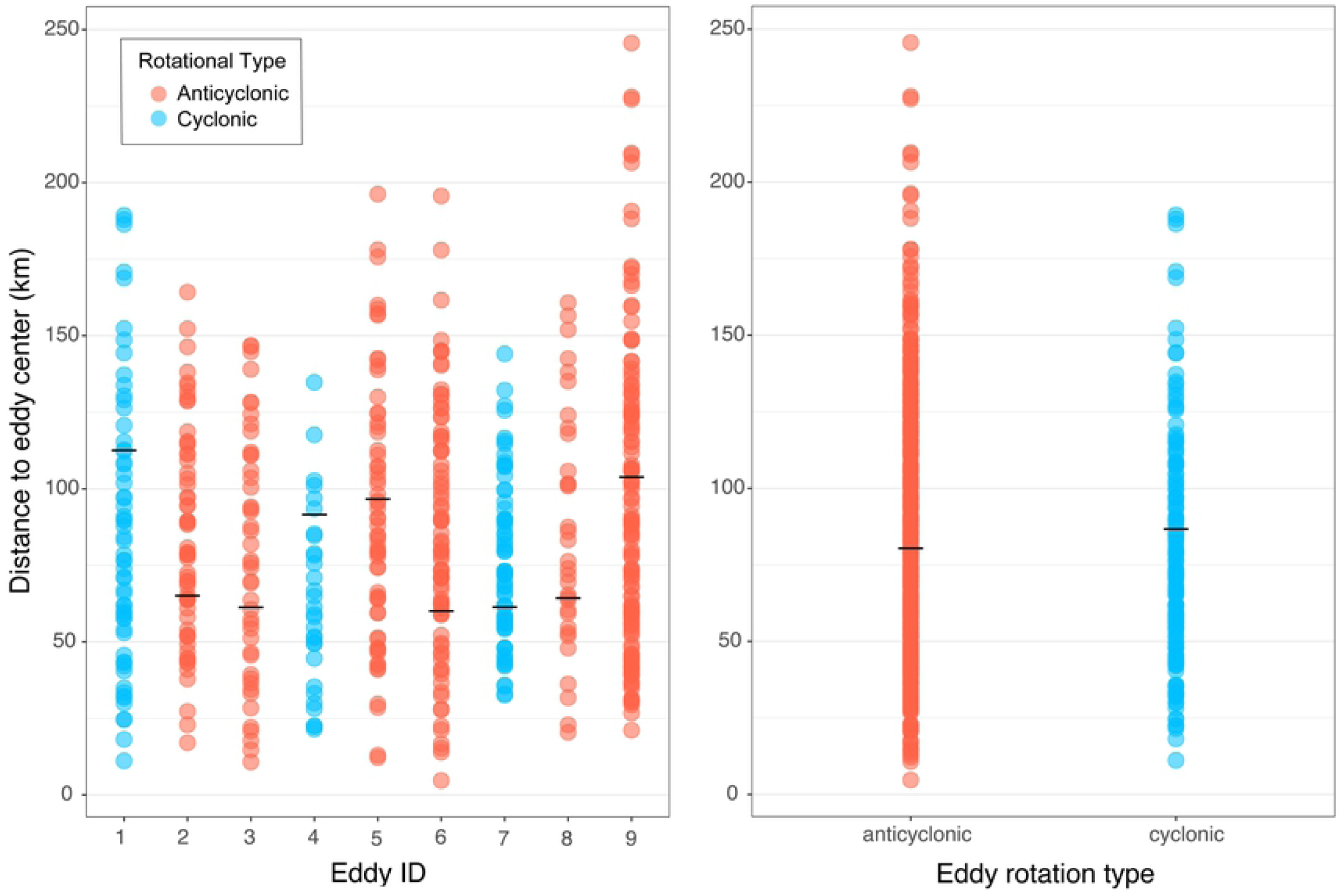
Distribution of plankton samples across eddies in relation to eddy core. Distance from plankton samples to eddy center (km) for (a) each eddy and for (b) each eddy rotational type. Anticyclonic eddies are in blue and cyclonic eddies are in red. The (a) radius or (b) mean radius is depicted with the horizontal black line within the distributions of points.

### Associated eddy and environmental data

Daily mesoscale eddy center location, diameter, rotational direction, and rotational speed were obtained from the Mesoscale Eddy Trajectory Atlas version 1.0, produced by SSALTO/DUACS and distributed by AVISO+. Distance from the eddy center to the nearest point of land was calculated in R with the gDistance function (‘rgeos’ package [27]). Coastlines were downloaded from NOAA (GSHHG data version 2.3.7; June 15, 2017). CPR location data were collocated with the eddy database in Python using the Numpy, Xarray, and Pandas libraries [28–30]. Only CPR data that were within mesoscale eddies or within 200 km from the outer edge of eddies were used within this study. Additionally, we excluded any eddies that were not transected by the CPR (i.e., eddies that were close to the CPR, but were not sampled both inside and on their periphery). Canadian Meterological Center (CMC) global Sea Surface Temperatures (SSTs) version 2.0 were collocated with remaining CPR sample points using linear interpolation in time and space [31]. To test the effect of eddy structuring on plankton abundance, standardized distances were calculated to the eddy edge from 1) within the eddy, by subtracting the eddy radius from the distance of the CPR sample to the eddy center divided by the radius, and 2) the outside the eddy, by subtracting the distance to the edge from the radius divided by the maximum sample distance from the edge (164 km). The distance of samples from the center of each eddy was visualized to ensure comparability of eddy distances to the edge between types (Fig 1). In total, 3 cyclonic and 6 anticyclonic eddies sampled twice across a period from 2002 to 2013 were used to describe eddies and for analyses (Fig 2).

**Fig 2.**
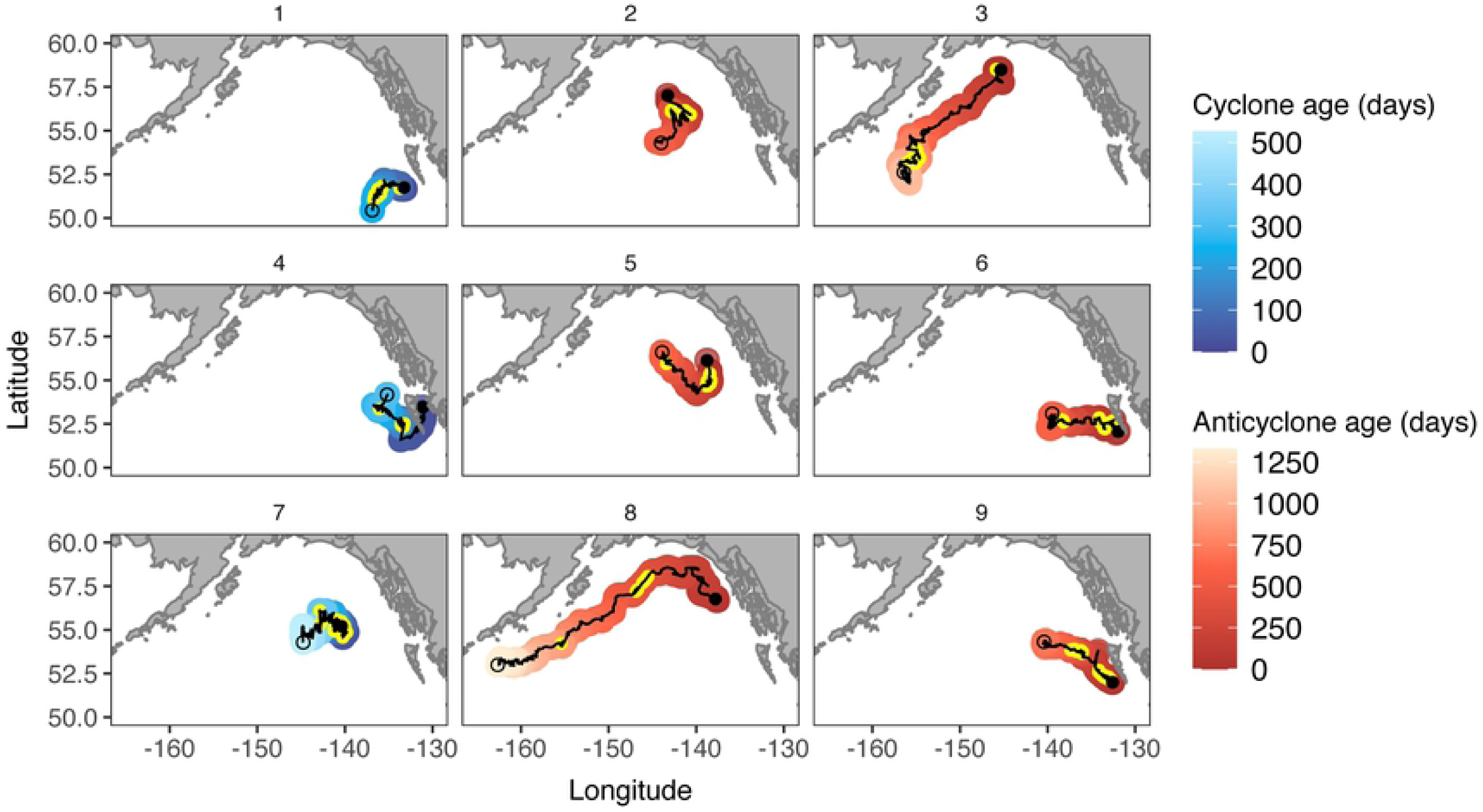
Eddy sampling locations in different regions and at different eddy ages. Sampling of cyclonic (red) and anticyclonic (blue) eddies within the Gulf of Alaska. Eddy paths begin with a solid black circle and end with an open black circle. Yellow circles depict sampling periods (both inside and within 164 km from the edge of the eddies). The age in days of the eddy, depicted in darker (younger) to lighter (older) hues, demonstrates the relative lifespans of the individual eddies.

### Statistical Analysis

Within each eddy rotational type (anticyclonic and cyclonic), the relative differences in mean log abundance between taxonomic groups were assessed. First, a nonparametric Kruskal-Wallace rank sum test (kruskal.test function; base R) was used to identify if significant differences (P<0.05) between groups were present. Where significant differences were found, Dunn’s test post hoc pairwise comparisons were made (DunnTest function; ‘DescTools’ package [32]) using a Bonferroni correction to determine which taxonomic groups were significantly different.

To determine how the physical dynamics of each eddy rotational type influenced community structure, the abundance of each plankton group was modeled in relation to the distance to the eddy edge. Generalized additive mixed models (GAMM; gam function; ‘mgcv’ package [33]) were used to visualize patterns in abundance across eddies. In each model, eddy rotational type (cyclonic or anticyclonic) was included as a factor interaction with distance to eddy edge to determine how relationships varied between rotational types. The following potentially influential predictors were also included in the models: eddy age as percentage of lifespan, eddy rotational speed, eddy distance to land, day of year, hour of day, eddy ID, and year. Sea surface temperature was not included as a predictor as it was highly correlated (>0.6) with day of year. Thin plate regression basis splines were used for all predictors except temporal predictors where cyclic cubic regression splines were used, and eddy ID and year where random effects were used. A tensor interaction term was included for time of day and day of year, which reduced the degrees of freedom but improved model residuals. In zooplankton models, total phytoplankton (diatoms + dinoflagellates) was also included as a predictor. Samples with outlier phytoplankton counts were removed (n=2). Models were checked for overfitting by comparing the R^2^ and deviance explained between a training and test set of data.

Where significant patterns across eddy types were found, the relationships were modeled again using only Haida eddies (i.e., eddies generated near the Haida Gwaii archipelago; n=301) to parse the effects of origin on the abundances of plankton across eddy type. Haida eddies were the only eddy type with enough samples in each eddy rotational group for meaningful analysis (Fig 1 and S1 Table). Next, to parse the effects of eddy life stage, the same relationships from all regions were modeled separately within life stage groups defined by an approximate early stage (<25% of lifespan; n = 277), mature stage (≥25% and ≤80% of lifespan, n=237), and a combined mature/decay stage (≥25% of lifespan, n = 290) [11]. We examined the mature phase combined with the decay phase because the sample size for mature stage cyclonic eddies was low (n=48). The random effect of year was removed from models separated by life stage. A quasi-Poisson family was used for all GAMs to account for over-dispersed count data as this distribution performed better than Tweedie or negative binomial distributions. To optimize smoothness selections, a restricted maximum likelihood method was used and an extra penalty was added to all terms. Models were checked for concurvity and nonrandom predictors (with the exception of the main effects of the interaction term) were removed where the observed and estimated values were >0.80). Model fits were examined by visually inspecting the model residuals and basis dimensions using the gam.check function [33].

The majority of copepod taxa are omnivorous and assumed to be heavy grazers of phytoplankton, therefore, to examine whether or not the effect of the distance to eddy edge on zooplankton was direct, or indirectly mediated by phytoplankton abundance, we tested the influence of total phytoplankton on total (large + small) copepods. We used causal mediation analysis (‘mediation’ package; [34]) with 1000 nonparametric bootstrapped simulations and bias-corrected confidence intervals on the GAMs selected as described above, but we used a Poisson family distribution. The mediator and outcome models were examined separately for each eddy type in order to remove any interaction with the ‘treatment’ variable (distance to eddy edge) and the tensor interaction term (hour by day of year) was also removed to adjust for the reduced sample size. The remaining covariate terms were kept consistent between the mediator and outcome models. Because the distance to eddy edge was continuous and because we were interested in direct vs indirect effects of phytoplankton within the eddy, a contrast was applied from the center of the eddy to the eddy edge (control = 0; treatment = -1). As an additional check, the data were filtered to remove all points outside the eddies and a manual stepwise mediation (as established in [35]) was applied to GAMs where distance to eddy edge and total phytoplankton were included without smoothing terms (i.e., as linear terms). The resulting linear parameter estimates for each pathway were then examined. All statistical analyses were performed in R version 3.6.3 [36].

## Results

### Origin and physical characteristics of cyclonic and anticyclonic eddies

Two anticyclonic eddies originated in the Haida region, two were from the Sitka region, and two were from the Alaska Stream region (regions as described in [37]). Two eddies (from the Alaska Stream region and Sitka region) traveled the greatest longitudinal distance westward across the Gulf of Alaska within the Alaska Stream region (Fig 1). As expected, not all eddies had characteristic relative core temperatures during the periods of CPR sampling (S1 Table; Sun et al., 2019): two anticyclonic eddies contained cooler cores than their surrounding water and the cyclonic eddies, and one cyclonic eddy contained a much warmer core than the surrounding waters (S1 Fig). General eddy characteristics of each eddy are summarized across the entire lifespan (S1 Table) and within the sampling periods (Table 1). All eddies contained samples from the intensification (10 to 25% lifespan) and mature stages (25 to 80% lifespan) with the exception of eddies 3 and 6 that were sampled in the formation stage (0–10% lifespan) instead of intensification stage and eddy 4 that was sampled in the decay stage (80–100% lifespan) instead of the mature stage (stages as defined in [38]; Table 1). All eddies were formed during the spring or early summer, but one anticyclonic eddy from the Haida region was formed in the fall (S1 Table). Compared to cyclonic eddies, the anticyclonic eddies had faster mean rotation speeds (22 cm s^-1^ vs. 9 cm s^-1^) and longer mean lifespans (2.2 years vs. 1 year; S1 Table).

### Relative abundance of plankton between taxonomic groups

Within both anticyclonic and cyclonic eddies, the diatoms had a significantly greater ranked mean abundance than all other groups (P<0.001) with the exception of large copepods in anticyclonic eddies, where there was no significant difference (P=0.33). After diatoms, large and small copepods were greater in mean abundance than all other groups (P<0.001) except for microzooplankton in cyclonic eddies only, where there was no significant difference (P=1.0). Microzooplankton were also relatively high in abundance and were significantly more abundant than all remaining groups in both eddy types (dinoflagellates, euphausiids, hyperiids, pteropods, chaetognaths, and larvaceans; P<0.001). The remaining groups were relatively equivalent in mean abundance with the exceptions that there were more hyperiids and fewer larvaceans than dinoflagellates in the anticyclonic eddies (P<0.01) and more chaetognaths than dinoflagellates, pteropods, larvaceans in the cyclonic eddies (P<0.01; S2 Fig). Due to the relatively poor CPR sampling of dinoflagellates, pteropods, and larvaceans, these taxonomic groups were excluded from further analysis.

### Phytoplankton and zooplankton structure across eddies

Among the phytoplankton taxonomic groups, the abundance of diatoms was lower near the center of anticyclonic eddies and higher near the edge and the surrounding waters of anticyclonic eddies (Fig 3). Though marginally insignificant, a similar pattern with more uncertainty around the predicted abundance was detected in cyclonic eddies. There were no significant patterns in dinoflagellate abundance across either eddy type. Among the zooplankton taxonomic groups, euphausiids, small copepods, and microzooplankton exhibited inverse abundance relationships to diatoms within anticyclonic eddies, such that their abundances were highest at the center of the eddy compared to the edge and surrounding waters. Conversely, within cyclonic eddies, euphausiids, small copepods, and microzooplankton were less abundant near the center of the eddies and increased in abundance toward the eddy edge. Large copepods and hyperiids were highest in abundance mid-distance between the eddy center and edge (hereafter the “radial midpoint”). Where significant patterns were observed, peak zooplankton abundances were within or on the edge of all eddies except for the hyperiid group, which reached a maximum in the surrounding waters. There were no significant patterns in chaetognath abundances across either eddy type, thus further analysis of this group was not explored. Model residuals generally met assumptions of normality; however, the small copepod model residuals exhibited a positive skew.

**Fig 3.**
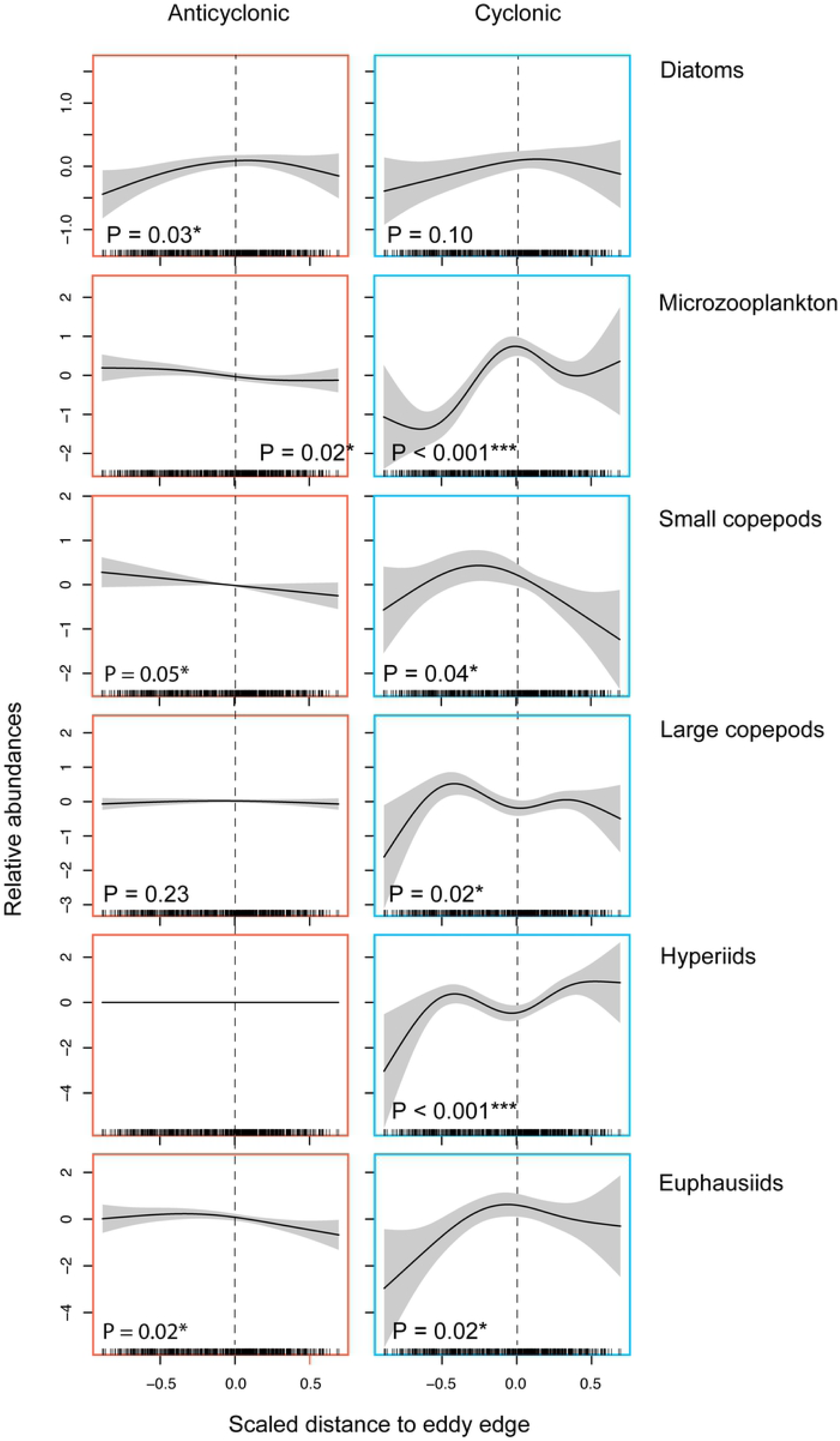
Patterns of plankton abundance across eddies vary by taxonomic group and eddy rotational type. Smoothed relative abundances of taxonomic groups relative to partial effects of eddy edge by eddy type (anticyclonic in red, cyclonic in blue) from GAMs. The y-axes for each model (i.e., taxonomic group) are set to the same scale for comparison across eddy type, but the scale varies between models. Distance to the eddy edge on the x-axis is scaled for all models such that -1 indicates samples closest to the center of the eddy, 0 indicates eddy edge (dashed line), and 1 indicates samples closest to the cutoff point (164 km) outside of the eddy. Groups where neither eddy type showed a significant relationship are not shown (dinoflagellates, larvaceans, and chaetognaths). Hash marks at bottom of plot indicate the sampling locations for the entire model.

Within Haida eddies only, similar patterns in plankton abundances across eddy types were observed but with notable exceptions. Across anticyclonic eddies, there were no significant patterns in diatom abundance and across cyclonic eddies there were no significant patterns in large copepod abundance (Fig 4). Within anticyclonic eddies, microzooplankton abundance was lower within the eddy than the surrounding waters, similar to the pattern in cyclonic eddies.

**Fig 4.**
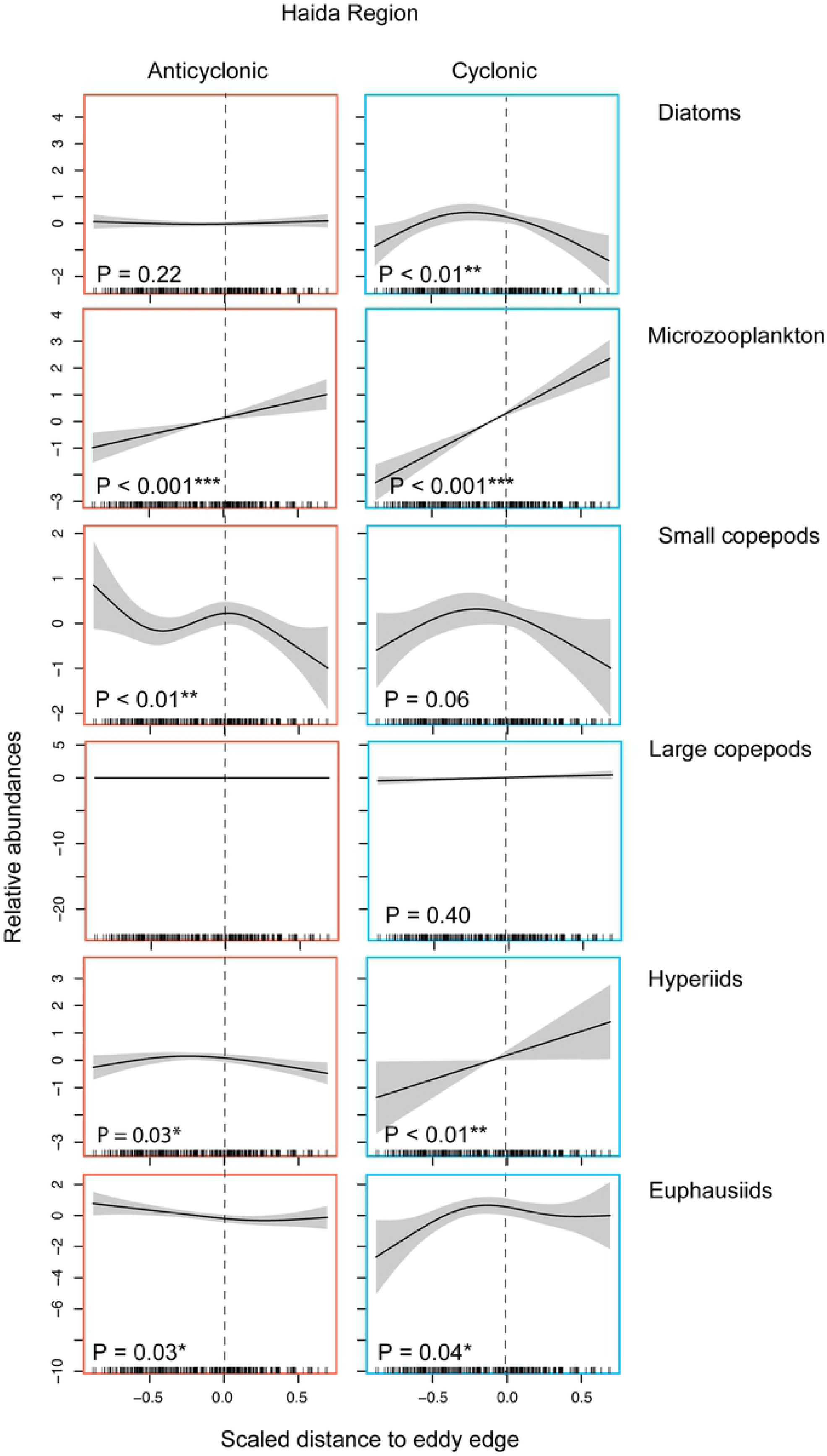
Patterns of plankton abundance across Haida eddies vary by taxonomic group and eddy rotational type. Smoothed relative abundances from GAMs of taxonomic groups relative to partial effects of eddy edge by eddy type (anticyclonic in red, cyclonic in blue) within eddies originating from the Haida region only. The y-axes for each model (i.e., taxonomic group) are set to the same scale and center smoothed for comparison across eddy type, but the scale varies between models. Distance to the eddy edge on the x-axis is scaled for all models such that -1 indicates samples closest to the center of the eddy, 0 indicates eddy edge (dashed line), and 1 indicates samples closest to the cutoff point (164 km) outside of the eddy. Hash marks at bottom of plot indicate the sampling locations for the entire model.

Additional spatial patterns in abundance across eddy types emerged when eddies were separated by life stage (see Fig 5 for following results). The anticyclonic eddy diatom pattern shifted from higher abundance near the eddy edge during the early stage to higher abundance at the radial midpoint during the mature stage. Among the zooplankton in anticyclonic eddies, the pattern of greater abundance at the eddy center disappeared for microzooplankton in all stages and for small copepods during the early stage. Small copepods were more abundant toward the eddy center in the mature and decay stages. Conversely, euphausiids were more abundant at the eddy center during the early stage but not the latter eddy life stages.

**Fig 5.**
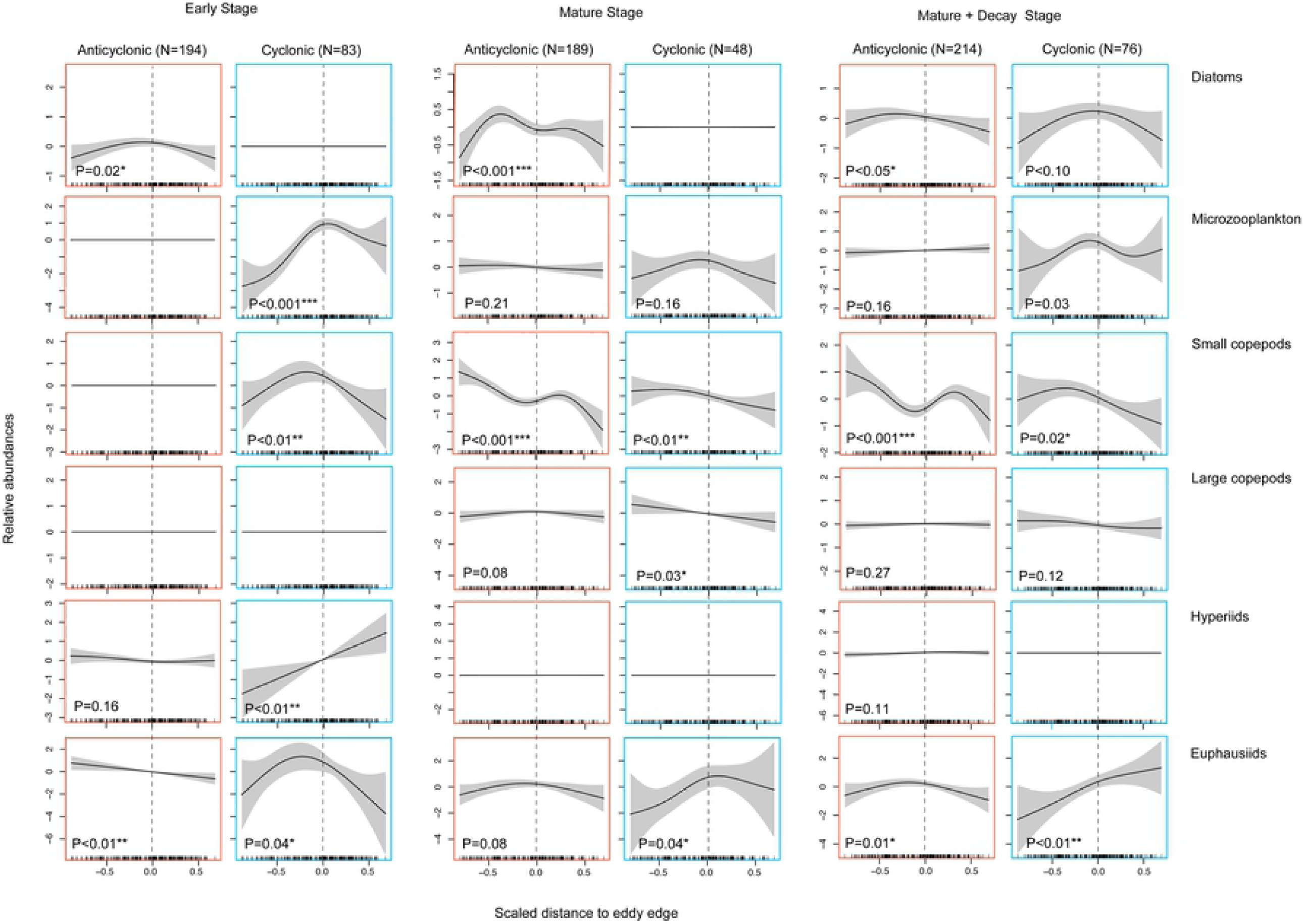
Patterns of plankton abundance across eddies vary by taxonomic group, eddy rotational type, and eddy life span. Smoothed relative abundances of taxonomic groups in the early stage, the mature stage, and the mature stage combined with the decay stage of the eddy lifespan relative to partial effects of eddy edge by eddy type (anticyclonic in red, cyclonic in blue) from GAMs. The y-axes within each model (i.e., taxonomic group) are set to the same scale for comparison across eddy type, but the scale varies between models and between life stages. Distance to the eddy edge on the x-axis is scaled for all models such that -1 indicates samples closest to the center of the eddy, 0 indicates eddy edge (dashed line), and 1 indicates samples closest to 164 km cutoff point outside of the eddy. Hash marks at bottom of plot indicate the sampling locations for the entire model.

Within cyclonic eddies, the marginal diatom abundance signal was penalized out of the model when split by stage, with the exception of the ‘mature + decay’ stage. When the mature stage included the decay phase the marginal pattern of peak diatom abundance near the eddy periphery emerged. Among the zooplankton in cyclonic eddies, abundance patterns during the early stage were similar to the full model (all stages), with the exception of large copepods. There was no significant pattern in large copepod abundance during the early stage and instead a pattern appeared in the mature stage but with a linear increase toward the cyclonic eddy center. Small copepods were less abundant at the eddy edge during the latter eddy life stages, but still higher in abundance within the eddies than surrounding waters. Their abundance near the eddy center, however, was still lower than in the anticyclonic eddies. Hyperiids exhibited no patterns in abundance across cyclonic eddies in the latter life stages. Euphausiids became more abundant in surrounding waters during the ‘mature + decay’ stage.

### Impact of phytoplankton on zooplankton

Full GAMs by taxonomic group indicated that, with the exception of euphausiids, each zooplankton taxonomic group had linear or curvilinear relationships with total phytoplankton abundance (S4 Fig). A positive linear relationship was detected for small copepods (and total copepods). All relationships became more variable with exponentially larger phytoplankton counts and microzooplankton increased in abundance until this point, when they then slightly decreased (S4 Fig). When models were parsed by rotation type for mediation, total copepod abundance initially decreased with increasing total plankton then increased when total phytoplankton reached counts of approximately 100,000 cells per sample in anticyclonic eddies (Fig 6). In cyclonic eddies, total copepod abundance asymptotically increased with total phytoplankton abundance (Fig 6). Within anticyclonic eddies, the bootstrapped mediation models did not show any significant effects, likely due to inconsistent mediation [39], where the mediation pathway suppresses the direct pathway due to opposing signs (or curvilinearity, in this case). The stepwise approach revealed that in the anticyclonic eddies, the pathways a and c’ (Fig 6a) were opposing in sign (estimates=-0.32 and 0.41, respectively), and that although the effect of total phytoplankton on total copepods (pathway b; Fig 6b) was significant (P<0.001), the effect size was negligible (estimate=0.00) and no mediation occurred. Within cyclonic eddies, the bootstrapped mediation models indicated no mediation (averaged: P=0.23) with only direct and total effects of phytoplankton on zooplankton (averaged direct estimate: -189, 95% CI [-43, - 275], P=0.03; total estimate=-219, 95% CI [-71, -310], P=0.01; respectively).

**Fig 6.**
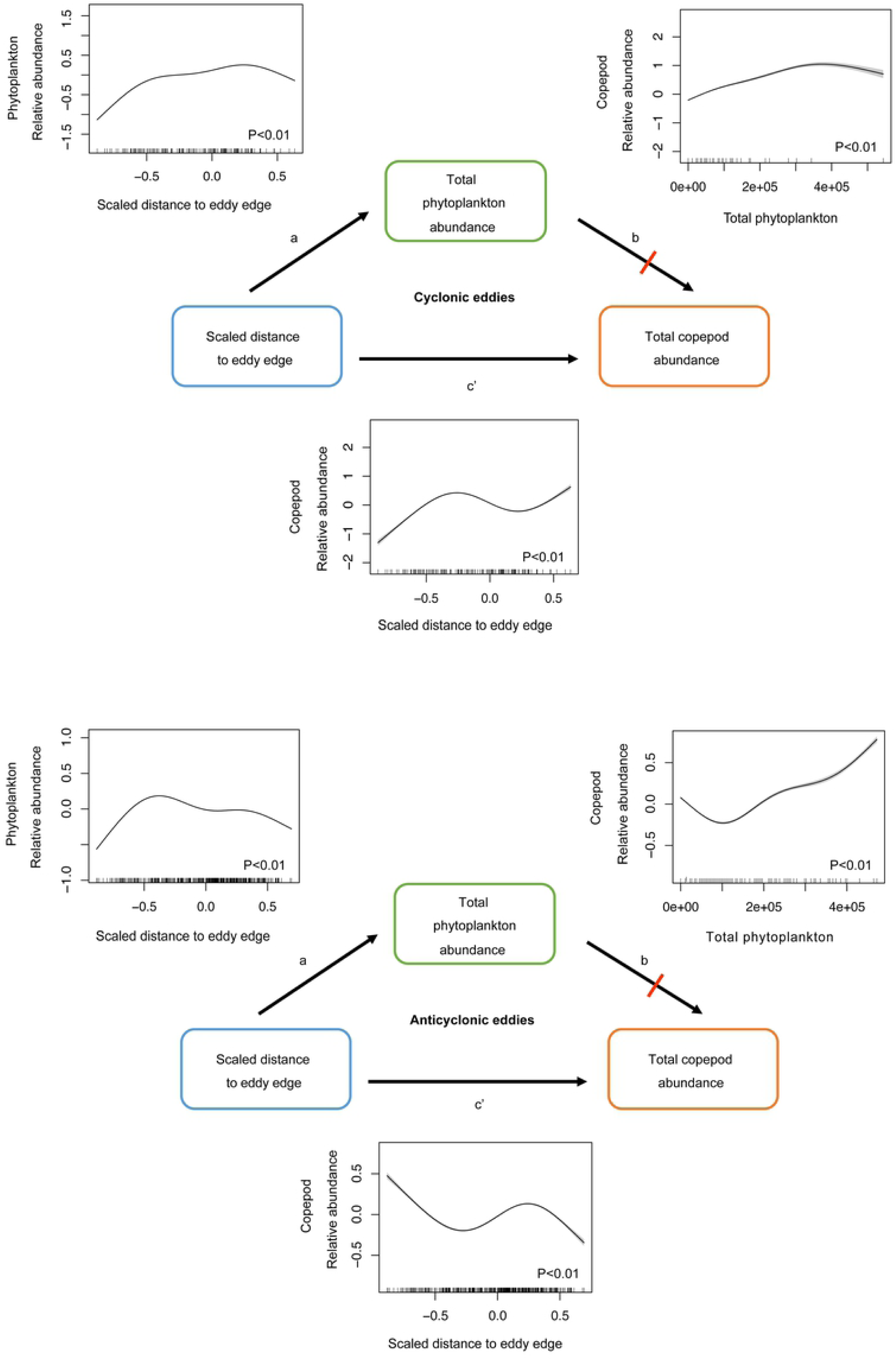
Distance to eddy edge directly affects zooplankton abundance. Anticyclonic (top) and cyclonic (bottom) eddy GAM estimates of relative abundance of total phytoplankton predicted by distance to eddy edge (pathway a) and relative abundance of total copepods predicted by total phytoplankton (pathway b) and distance to eddy edge (pathway c’). Red line indicates mediation does not occur. Additional covariates (hour, day of year, and random effects of eddy ID and year) are not shown. Mediation pathways are represented by a and b and the direct pathway is represented by c’. In the bootstrapped mediation, the mediator model tests pathway a and the outcome model simultaneously tests the effects c’ while controlling for the effect of b.

## Discussion

In this study we gain a more complete and consistent description of how eddy dynamics associated with rotational type, origin, and life span affect the structure of plankton communities in the pelagic Gulf of Alaska using high resolution, *in situ* CPR data inside and outside several eddies. Phytoplankton abundances were highest near eddy edges for both eddy rotational types (Figs 1 and 6), consistent with satellite observations [38], whereas zooplankton abundance was higher near the edge of cyclonic eddies but higher near the center of anticyclonic eddies. These overall plankton community patterns were observed within eddies originating from the same region (i.e., the Haida region) for small copepods, hyperiids, and euphausiids, indicating that differences for these groups were largely due to rotation type (and possibly other eddy dynamics), not origin region. However, for microzooplankton and large copepods, origin region may have been important. Moreover, eddy age was a determining factor in plankton abundance within eddies, with some abundance patterns only found in the early stage (e.g., hyperiids) or only in the mature stage (e.g., large copepods), and with some abundances shifting from the center to the edge of eddies with time (e.g., euphausiids). Finally, when explored within the copepod group, the relationship between the distance to the eddy edge and abundance was not mediated by phytoplankton abundance.

### Various eddy dynamics determine plankton composition and organization

In our study, varied contributions of complex eddy hydrodynamics and associated biogeochemistry could explain the similar phytoplankton abundance patterns observed in both eddy types. Typically, nutrients (e.g., iron, which may be limiting offshore) and nutrient-dependent phytoplankton are vertically pumped in opposing patterns for each rotational type [13]. During formation, anticyclonic eddy pumping forces vertical upwelling on the outer edges of the eddy and convergence and downwelling at the center; these characteristics are consistent with warm, positive sea-level anomalies, and negative chlorophyll anomalies in the eddy core [40–42]. Opposite dynamics and patterns occur during cyclonic eddy formation [9,41,42]. The North Pacific, however, is known to have anticyclonic cold core and cyclonic warm core eddies [43], in addition to chlorophyll rings and high biomass around the peripheries of both eddy types [14,38]. Argo floats have been used to identify that radial displacement (i.e., displacement along the radius from the eddy center) and edge convergence occurs in anticyclonic eddies even when wind-driven eddy-induced Ekman pumping that causes inverse upwelling patterns to those at eddy intensification could enhance chlorophyll at the eddy center [8,10,44]. Thus, in our study, the peripheral signal of diatoms observed in both eddy rotational types may be attributable to strong vorticity of the anticyclonic eddies. Cyclonic eddies are much slower and therefore less likely to be affected by weaker Ekman pumping than internal eddy dynamics; however, Ekman pumping could still cause downwelling at the eddy center and divergence and upwelling at the eddy edge near the surface [45], consistent with our observation.

### Eddy origin and life stage influences on phytoplankton

Further examination of eddies by their origin region and life stage illuminated the importance of these respective influences on the composition and organization of plankton communities. For example, early-stage anticyclonic eddies in the Haida region showed no pattern in diatom abundance across eddy edges in contrast to our model with all eddies combined; suggesting that the pattern of higher edge abundance in the latter model may be influenced by eddy formation in more northerly latitudes. Moreover, long-lived eddies (3 and 8), sampled while propagating in the Alaska Stream (Fig 1), had faster rotational speeds relative to other eddies (S1 Table); thus, both Ekman pumping and the radial displacement of entrapped diatoms could account for the observed spatial pattern. Diatoms are more abundant in the northern Gulf of Alaska and along the Alaska Peninsula [46], which could support diatoms near the periphery through eddy stirring throughout the anticyclonic eddy lifespan. Indeed, the mature stage anticyclonic eddies exhibit high abundance near the radial midpoint, consistent with satellite observations of chlorophyll anomaly dipoles that can be induced by the horizontal stirring of water masses [4,11].

In contrast, diatoms within cyclonic eddies from the Haida region showed peaks in abundance near the eddy edge. This pattern disappeared when the eddy data were parsed into early and mature stages, but when the decay phase was included (i.e., mature and decay phase combined), peak diatom abundance unexpectedly appeared near the eddy edge, but was not significant. This diatom structuring across eddy life stages indicates that a combination of regional influence, decay phase eddy pumping, and Ekman pumping of nutrients at the cyclonic eddy periphery supported diatom abundance. The proliferation of diatoms in the cyclonic Haida eddies may be due to greater nutrients in the water mass at formation, as eddy 4 formed between Haida Gwaii and the coast of British Columbia. The lack of diatom patterns in anticyclonic Haida eddies may have been confounded by an iron deposition experiment in July 2012 when 100 tons of iron sulfite were dumped into the ocean 300 km off the west coast of Haida Gwaii; this action resulted in higher-than-average zooplankton abundance that could have increased grazing pressure on diatoms [47]. In summary, the overall influence of eddies on phytoplankton distribution within and across eddies is best understood when accounting for regional water mass characteristics and eddy life stage.

### Eddy origin and life stage influences on zooplankton

Likewise, among the zooplankton, examination of abundance variation by origin region and eddy life stage was revealing. Contrary to our hypothesis that early life stage dynamics would have a greater influence on community organization, in anticyclonic eddies, only euphausiids exhibited peak abundances near the eddy center in the early life stages with their abundance becoming lower at the center relative to the eddy edge in later eddy life stages. Early-staged anticyclonic eddies are known to form with warmer, more oligotrophic waters entrapped at their cores [13]. Consistent with the entrapment of less productive waters, there was no pattern of abundance in the other zooplankton taxonomic groups in the early eddy life stage. Small copepods became more abundant near the center of mature and decay stage anticyclonic eddies when ageostrophic perturbations and wind-driven Ekman pumping could lead to upwelling at the anticyclonic eddy centers [6].

As expected, the cyclonic eddies in this study became less “productive” as they aged and traversed the Gulf of Alaska relative to anticyclonic eddies. Slower-rotating cyclonic eddies should also be expected to have weaker upwelling of nutrients and thus become depleted more rapidly, and with their shorter overall lifespans the regeneration of zooplankton communities is less likely. Indeed, the taxonomic group with more mobility (i.e., euphausiids) were higher in abundance outside of the cyclonic eddies during the decay stage. The relatively greater abundance of large copepods sustained in the cyclonic eddy core relative to surrounding waters in later stages (a pattern that contrasted with the other groups) may be more reflective of a lack of offshore large copepods, thus highlighting the potential importance of cyclonic eddies in horizontally transporting large copepod species in the Gulf of Alaska.

### Biological structuring of zooplankton

In addition to the effects of hydrodynamics, zooplankton structuring across eddies could have been influenced by their biological and life history traits. For example, where smaller zooplankton may be more susceptible to physical forcing, such as radial displacement, larger zooplankton can swim vertically to remain stable in the water column [48]. Diel vertical migration is thought to allow some species to avoid mixing dynamics that force the horizontal export or “leaking” of zooplankton from eddies [49]. Large copepods in this study, with the exception of those within mature cyclonic eddies, were uniform in structure across eddies similar to those sampled in [49], which was attributed to seasonal breeding cycles, during which spawn enter the eddy from below and remain beneath the mixed layer until vertical migration in the late summer. Correspondingly, most of our samples were from late spring/early summer. Hyperiids will also remain below the mixed layer during both day and night, feeding on copepods and other zooplankton [50], which could account for a similar uniform structure across later eddy life stages with some exceptions that could be attributed to differences in the movement patterns of gelatinous zooplankton hosts used by hyperiids [51]. Notably, small copepods in anticyclonic eddies persisted at the eddy core relative to the eddy edge in later life stages when other zooplankton like large copepods and hyperiids exhibited declines in abundance. Small copepods may have a preference or higher tolerance for warmer waters (i.e., warm core anticyclonic eddies), as they are more abundant in warmer temperatures compared to large copepods [52] and are known to concentrate at shallower depths in the mixed layer [53]. Small copepods also have shorter reproductive cycles, thus recruitment within longer-lived eddies and subsequent increases in small copepod abundance relative to large copepod abundance is more likely. When eddy pumping reverses in the later eddy life stages, small copepods may remain at the eddy center where there is greater upwelling rather than converging at the eddy edge due to vertical migration enabling them to escape radial displacement and predation by the larger carnivorous zooplankton.

The effect of grazers on controlling phytoplankton abundance, or conversely, the bottom-up effect of phytoplankton abundance on the location of grazers, complicates the assessment of physical structuring of plankton biological communities. To address this issue, we examined relationships between copepods (small and large) and phytoplankton (dinoflagellates and diatoms). We found that copepod distribution across eddies did not depend on diatom abundance. Some copepods may feed on other undetected phytoplankton species (i.e., those without hard shells or too small to be adequately retained by the CPR) and/or copepods may also consume microzooplankton at high rates (e.g., [54]). Other consumers of diatoms and/or copepods could potentially confound simple mediation pathways, especially considering that microzooplankton are also heavy grazers of diatoms in the Gulf of Alaska [55]. Alternatively, the physical forcing of the eddy hydrodynamics such as radial displacement and convergence that structures phytoplankton across an eddy may similarly force an overlap of zooplankton that is advantageous for the grazers. Depending on the life stage of the eddy, the overlap may also be determined by the spatial distributions of the organisms during entrainment by the eddy, which seems likely given the difference in pattern between phytoplankton and zooplankton abundance in the early stage. In support of a primary influence of eddy mechanics on distributions, Schmid et al. [24] found passive spatial structuring of larval fishes within an eddy in relation to their *Oithona* spp. copepod prey that they attributed to eddy structural mechanics. Eddy mechanics thus appear to play a key role in directly structuring low-motility zooplankton communities, which should indirectly structure more mobile, higher-level consumers such as krill, fish, seabirds, and marine mammals.

## Conclusion

Although we used a large number of CPR samples, there were few eddies available for investigating plankton communities both within and outside eddy margins. Fewer eddies lead to a wider margin of uncertainty within models for cyclonic eddies that were not as abundant in our study (n=3, Table 1). There is also greater uncertainty in the relative abundances of plankton near the center of cyclonic eddies as the CPR sampled farther from those eddy centers compared to anticyclonic eddies (Fig 2), but the patterns across both eddy types are still clear. We also note that the influence of eddies in shaping marine biological communities extends below the depth at which the CPR sampled (e.g.,[56]). Furthermore, it is known that other characteristics, such as rotational speed and specific eddy age, may influence associated planktonic communities [49,57], but we removed characteristics that exhibited high collinearity with other fixed variables in our model. Typically, from eddy formation and intensification to maturation and eventual decay, there is a complex spatiotemporal evolution of chlorophyll anomalies associated with eddies, relating to shifts in internal eddy pumping and peripheral stirring with surrounding waters [11]. However, day of year captures some of the variability associated with eddy aging, as well as temperature effects that differ with season. Despite these potential caveats, we were able to quantify the influence of major eddy characteristics on plankton communities by separating the data into early, mature, and decay life stages.

Cyclonic and anticyclonic eddies play important roles in directly structuring phytoplankton and zooplankton communities in the Gulf of Alaska, with their relative influences depending on formation region, eddy trajectory (e.g., where mixing occurs), and pumping dynamics that shift across the eddy lifespan. The mechanisms that structure plankton communities are undoubtedly multifaceted and include horizontal stirring, eddy pumping, and eddy-induced Ekman pumping. These processes distribute plankton in varied ways. Long-lived anticyclonic eddies with high rotational vorticity were important for stirring nutrients and sustaining productive phytoplankton and small copepod communities across the northern continental margin. This finding is in contrast to studies that show a significant reduction in nutrients in eddies after the first 4 months [17,58], but similar to other studies demonstrating the importance of anticyclonic eddies to marine ecosystems in other ocean basins [13,46,59,60]. As the cyclonic eddies decayed, they became relatively less productive than anticyclonic eddies for small copepods, but continued to be regionally important for the offshore transport of large copepods. Anticyclonic and cyclonic eddies provide differing ecosystem structuring to the Gulf of Alaska, shaping marine communities regionally based on the organisms that are encapsulated, the enhancement of production through eddy dynamics, and the stirring of plankton into surrounding waters.

## Datasets

The Mesoscale Eddy Trajectory Atlas products were produced by SSALTO/DUACS and distributed by AVISO+ (https://www.aviso.altimetry.fr/) with support from CNES, in collaboration with Oregon State University with support from NASA. CMC SSTs version 2.0 are available from the JPL PO.DAAC dataset identification CMC0.2deg-CMC-L4-GLOB-v2.0.

## Acknowledgements

We thank Dr. Brian Hoover for thoughtful discussions during data analysis and for comment on the first draft of this manuscript. We also thank the volunteer vessels and crews that towed the CPR devices and the members of the CPR program that processed the taxonomic data used in our study.

## Supporting information captions

**Supplemental Fig 1. Relative smoothed SST across each sampled eddy**. The estimated SST on the y-axis is from a generalized additive model with a centered factorial smoother (by ‘Eddy ID’) and a random effect of year. Anticyclonic eddies are in red, cyclonic in blue. Distance to eddy edge is scaled such -1 is the center, zero is the eddy edge, and 1 is the 150 km outside the eddy.

**Supplemental Fig 2. Relative abundance of taxonomic groups in each eddy rotational type**. Median log density and boxed interquartile range (IQR) for each taxonomic group for anticyclonic eddies (top) and cyclonic eddies (bottom). Whiskers indicate 1.5 x IQR and outliers are depicted as closed circles. Box plots for each taxonomic group are in descending order based on mean log density.

**Supplemental Fig 3. Full model partial plots and residual plots for each taxonomic group**. (A) Full generalized additive model fitted plots with all covariate partial effects. The y-axis shows relative log abundance scaled the same for each plot and is center smoothed – estimated degrees of freedom are in parentheses. (B) Plots for visual inspection of model assumptions: (top left) quantile-quantile plot of deviance residuals, (top right) histogram of residuals, (bottom left) residuals vs. linear predictor, (bottom right) response vs. fitted value.

**Supplemental Table 1. Summarized characteristics across eddy lifespan.**

**Supplemental Table 2. Generalized additive model results for each taxonomic group**.

